# SEX-DETector: a probabilistic approach to uncover sex chromosomes in non-model organisms

**DOI:** 10.1101/023358

**Authors:** Aline Muyle, Jos Käfer, Niklaus Zemp, Sylvain Mousset, Franck Picard, Gabriel AB Marais

## Abstract

**Data deposition:** During the review process, the SEX-DETector galaxy workflow and associated test datasets are made available on the public `galaxy.prabi.fr` server. The data as well as the tool interface are visible to anonymous users, but to use them, you should register for an account (“user Register”), and import the data library “SEX-DETector” (“Shared Data Data Libraries”) into your history. More instructions can be found in the “readme” file in this library. The user manual for SEX-DETector is available here: https://lbbe.univ-lyon1.fr/Download-5251.html?lang=en.

Paper submitted as a Genome Resource.

We propose a probabilistic framework to infer autosomal and sex-linked genes from RNA-seq data of a cross for any sex chromosome type (XY, ZW, UV). Sex chromosomes (especially the nonrecombining and repeat-dense Y, W, U and V) are notoriously difficult to sequence. Strategies have been developed to obtain partially assembled sex chromosome sequences. However, most of them remain difficult to apply to numerous non-model organisms, either because they require a reference genome, or because they are designed for evolutionarily old systems. Sequencing a cross (parents and progeny) by RNA-seq to study the segregation of alleles and infer sex-linked genes is a cost-efficient strategy, which also provides expression level estimates. However, the lack of a proper statistical framework has limited a broader application of this approach. Tests on empirical data show that our method identifies many more sex-linked genes than existing pipelines, while making reliable inferences for downstream analyses. Simulations suggest few individuals are needed for optimal results. For species with unknown sex-determination system, the method can assess the presence and type (XY versus ZW) of sex chromosomes through a model comparison strategy. The method is particularly well optimised for sex chomosomes of young or intermediate age, which are expected in thousands of yet unstudied lineages. Any organism, including non-model ones for which nothing is known a priori, that can be bred in the lab, is suitable for our method. SEX-DETector is made freely available to the community through a Galaxy workflow.

## Introduction

Species with separate sexes (males and females) represent ~95% of animals (Weeks, 2012). In angiosperms, although rarer, separated sexes (dioecy) can be found in ~15,000 species (Renner, 2014). ~20% of the crops (e.g. papaya, grapevine, strawberries, kiwi, spinach) are dioecious or derive from a dioecious progenitor (Ming et al., 2011). However, the mechanisms for sex determination remain unknown for most plant species and a number of animal species (Bachtrog, Mank, et al., 2014). In numerous cases, it is even unknown whether sex chromosomes are present. In angiosperms, dioecy has evolved from an ancestral hermaphrodite state 871 to 5000 times independently (Renner, 2014), but less than 40 sex chromosomes have been reported so far (Ming et al., 2011). This suggests that sex determination is unknown in 95 to 99% of the dioecious angiosperm species. The situation is even worse in other plants where only a handful of sex chromosomes have been described among the ~6000 dioecious liverworts, ~7250 dioecious leafy mosses and ~381 dioecious gymnosperms (Ming et al., 2011). Precise estimates of the frequency of dioecy in brown and green algae are currently missing and very few sex chromosomes have been described in those groups. Consequently, further research is required to describe the diversity of sex determination and sex chromosomes in non-model organisms (Bachtrog, Mank, et al., 2014).

Sex chromosomes were originally a normal pair of autosomes that, after acquiring sex-determining genes, stopped recombining and diverged from one another (Bachtrog, 2013). In male heterogametic systems, males are XY and females XX whereas in female heterogametic systems, females are ZW and males ZZ. In species with a haplodiploid life cycle, sex can be expressed at the haploid phase with U females and V males and diploid individuals are heterogametic UV (Bachtrog, Kirkpatrick, et al., 2011). Y, W, U and V chromosomes all have a non-recombining region that can be small or spread to most of the chromosome. Sex chromosomes with a small non-recombining region are homomorphic (X and Y of similar size), which makes their identification through cytology difficult. And yet this type of sex chromosomes is probably frequent in groups such as angiosperms where many dioecious species have evolved recently and must have young sex chromosomes (Ming et al., 2011), in groups such as fish where sex chromosome turn over is high (Mank and Avise, 2009), or in groups such as amphibians where occasional recombination limits sex chromosome divergence (Stock et al., 2013). In such cases, sequences are required to identify sex chromosomes.

However, obtaining well assembled sequences of sex chromosomes is extremely difficult due to the repeats that accumulate in their non-recombining regions (Charlesworth et al., 1994; Gaut et al., 2007). Only the costly use of BAC clones organised in a physical map makes it possible to completely assemble DNA sequences from non-recombining regions. This is why only a handful of sex chromosomes have been fully sequenced to date (<15), many of which have small non-recombining regions: the liverwort *Marchantia* (Yamato et al., 2007), the fish medaka (Kondo et al., 2006), the green alga Volvox (Ferris et al., 2010), the tree papaya (Wang et al., 2012) and the brown alga *Ectocarpus* (Ahmed et al., 2014). Producing high-quality assemblies is not always necessary and alternative, less expensive strategies have been recently developed for identifying sex chromosome sequences based on next-generation sequencing (NGS) data (reviewed in Muyle, Shearn, et al., n.d.).

A first category of approaches relies on the comparison of female and male DNA-seq (DNA sequencing using NGS) data (Akagi et al., 2014; Carvalho and Clark, 2013; Cortez et al., 2014; Vicoso and Bachtrog, 2011; Vicoso, Emerson, et al., 2013; Vicoso, Kaiser, et al., 2013). As these methods require a reference genome (either from the studied species or a close relative), they will be difficult to apply to non-model organisms because reference genomes are lacking and/or genomes are large and complex. Another method uses the ploidy of SNPs in order to identify sex chromosome sequences (Gautier, 2014) but requires the sequencing of a hundred individuals, which, depending on the sequencing method, could be too expensive for non-model organisms. The methods cited thus far require that X and Y sequences be divergent enough not to coassemble or map onto one another. This means that they will work well in old systems but will probably fail with recently evolved sex chromosomes. Other approaches work well on young sex chromosomes, such as the use of sex markers (inferred from polymorphism data or genetic maps) to identify scaffolds belonging to sex chromosomes in a genome assembly (Al-Dous et al., 2011; Hou et al., 2015; Picq et al., 2014). However, the need for a reference genome can again be a hindrance for many non-model organisms, especially those with large genomes. In such cases, studying the transcriptome rather than the genome can be a cost saving measure. RNA-seq gives direct access to gene sequences and their expression levels, which can be valuable for various biological analyses. Identifying which genes in a transcriptome are sex-linked (i.e. located on the non-recombining region of sex chromosomes) can be done through the sequencing of males and females and the analysis of SNPs. For instance, brothers and sisters from an inbred line can be sequenced and used to infer sex-linked genes (Muyle, Zemp, et al., 2012). But inbred lines are unlikely to be available in most non-model organisms. A strategy relying on the sequencing of a cross (parents and progeny of each sex) by RNA-seq has proved very successful in the identification of sex-linked genes through the study of allele segregation (see Figure 1). This strategy requires that X and Y copies coassemble and map onto one another in order to identify X/Y genes using X/Y SNPs (Figure 1.b). X copies can also be identified on their own if the Y copy is absent because of degeneration or if X and Y were too diverged to coassemble (Figure 1.c). Therefore, this strategy is better suited for sex chromosomes that have either a low or intermediate level of divergence. However, it will still provide X copies of sex-linked genes in old systems, as long as appropriate SNPs are present in the dataset (Figure 1.c). Crosses can be obtained in any organism that can be grown in controlled conditions and are a common resource since they are needed for building genetic maps. Hundreds of sex-linked genes were identified this way in species with unavailable fully sequenced genomes such as Silene latifolia (Bergero and Charlesworth, 2011; Chibalina and Filatov, 2011) or in species without any genomic resource such as *Rumex hastatulus* (Hough et al., 2014) and *Rumex acetosa* (Michalovova et al., 2015).

**Figure 1.**
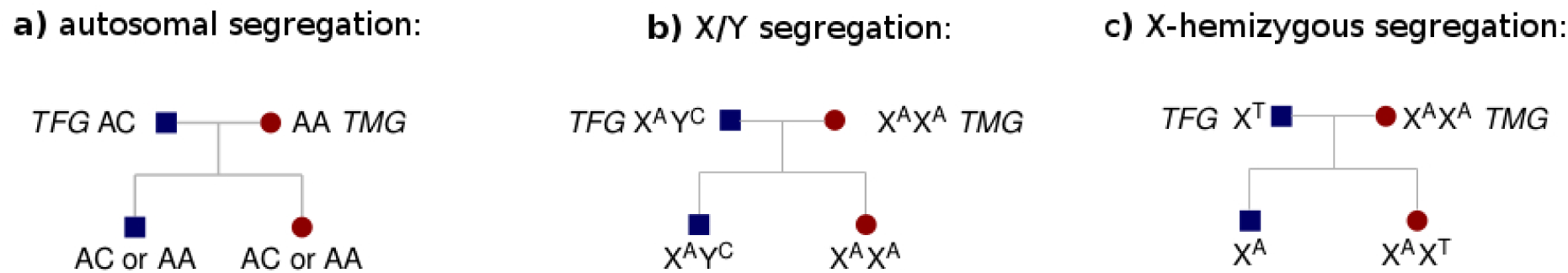
Examples for the three segregation types. a) autosomal, b) X/Y and c) X-hemizygous (when the Y copy was lost or was assembled in a separate contig). TFG stands for true heterogametic parent genotype and TMG for true homogametic parent genotype.

Although this RNA-seq cross-based strategy is very promising for studying sex chromosomes in nonmodel organisms, the existing approaches have a number of limitations due to the fact that inference of sex-linkage was done with empirical filters and without a statistical framework. Once RNA-seq reads have been mapped to a reference transcriptome, individuals need to be genotyped in order to study allele segregation in the cross (Figure 1). Genotyping was either done by filtering the number of reads at each locus with fixed thresholds (Bergero and Charlesworth, 2011; Chibalina and Filatov, 2011) or by using genotypers designed for DNA-seq data (Hough et al., 2014; Michalovova et al., 2015). This is problematic in the case of sex-linked genes where the Y allele is frequently less expressed than the X (reviewed in Bachtrog, 2013) and can be confounded with a sequencing error in RNA-seq data. Read number thresholds determined empirically for a given dataset can be sub-optimal for another dataset with different sequencing depth. In many cases, progeny individuals were pooled separately for each sex for sequencing in order to lower costs (Bergero and Charlesworth, 2011; Chibalina and Filatov, 2011; Michalovova et al., 2015). However, this makes it impossible to differentiate between all individuals of the pool or only a few being heterozygous, a criteria that is crucial to disentangle sex-linked genes from autosomal ones (Figure 1). Finally, sex-linked genes were filtered for having more than a given number of sex-linked SNPs (Chibalina and Filatov, 2011; Hough et al., 2014), and for not having any autosomal SNPs (Bergero and Charlesworth, 2011; Michalovova et al., 2015). These arbitrary filters clearly limit the application of this strategy to any dataset and probably prevent the detection of many true sex-linked genes. Also, a method allowing the study of UV systems is currently lacking.

Here, we propose a probabilistic method called SEX-DETector that solves the caveats of previous RNA-seq cross-based approaches and works on any sex chromosome type (XY, ZW, UV). The method was designed to discover as many sex-linked genes as possible from the data, while keeping inferences extremely reliable for downstream biological analyses. The pipeline, implemented in a galaxy workflow, was tested on empirical and simulated data and proved very promising for the discovery of many sex chromosomes and sex-linked genes in non-model organisms, especially in young systems.

## Materials and Methods

### The probabilistic model

#### Observed and hidden data

The data consist of genotypes in a cross (parents and progeny of each sex), at each position of every contig and can typically be obtained from RNA-seq experiments. The model aims to describe the transmission of alleles from parents to progeny, in the given cross, in order to infer whether a gene is sex-linked i.e. if it is located in the non-recombining region of sex-chromosomes (see Figure 1). The observed data, denoted by *G*, consist of the observed genotypes of the parents and the progeny, and we suppose that the probabilities of observing these genotypes depend on unknown information or hidden variables that we want to recover:

**The segregation type *S*** describes whether the studied locus is autosomal or sex-linked, which influences allele transmission from parents to progeny. There are three segregation types *j*: autosomal (*j* = 1), X/Y (or Z/W, *j* = 2) and X-hemizygous when the Y allele is absent (or Z-hemizygous when the W allele is absent, *j* = 3). The probability of segregation type *j* for position *t* in contig *k* is ℙ(S_*ktj*_ = 1) = *π_j_*.
**The true homogametic parent genotype TMG** (for true mother genotype in the case of male heterogamety) is introduced to account for genotyping errors that can cause the observed genotype to differ from the true genotype. There are ten possible genotypes *m* for the homogametic parent: `AA`, `AC`, `AG`, `AT`, `CC`, `CG`, `CT`, `GG`, `GT` and `TT`. The probability for the true homogametic parent genotype of being *m* at position *t* of contig *k*, given segregation type *j* is: 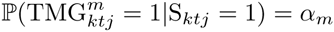. It is assumed that the true mother genotype frequencies do not differ between autosomal and sex-linked loci, so that parameter α is independent from segregation type *j*.
**The true heterogametic parent genotype TFG** (for true father genotype in the case of male heterogamety). The possible genotypes n for the heterogametic parent depend on the segregation type of the studied locus: there are ten possibilities for an autosomal segregation type (`AA`, `AC`, `AG`, `AT`, `CC`, `CG`, `CT`, `GG`, `GT` and `TT`), twelve for an X/Y (or Z/W) segregation type (`X`^A^`Y`^C^, `X`^C^`Y`^A^, `X`^A^`Y`^G^, `X`^G^`Y`^A^, `X`^A^`Y`^T^, `X`^T^`Y`^A^, `X`^C^`Y`^G^, `X`^G^`Y`^C^, `X`^C^`Y`^T^, `X`^T^`Y`^C^, `X`^G^`Y`^T^, `X`^T^`Y`^G^) and four for an X-hemizygous (or Z-hemizygous) segregation type (`X`^A^, `X`^C^, `X`^G^, `X`^T^). Given the segregation type *j*, the probability for the true heterogametic parent genotype of being *n* at position *t* of contig *k*, is: 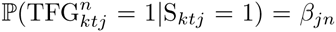. It is also assumed that the true parental genotype frequencies do not differ for autosomal loci (*β*_1_ = *α*).
**Genotyping error GE:** this variable describes whether there has been a genotyping error made for the studied individual *i*. It depends on the segregation type and the true parental genotypes of the studied locus. The probability for a genotyping error of having occured for individual *i* at position *t* in contig *k*, given the segregation type *j* and the true parental genotypes *m* and *n* is: 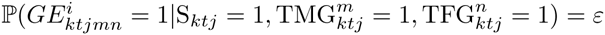.
**Y (or W) genotyping error YGE:** this variable accounts for the fact that genotyping errors are more common for Y and W alleles due to degeneration and lower expression. A Y or W genotyping error can only occur in a heterogametic individual *i*_het_ and in a X/Y segregation type (*j* = 2). Similarly to the genotyping error GE, it depends on the true parental genotypes. The probability for a Y or W genotyping error of having occured for individual *i* of sex *r* at position *t* in contig *k*, given the segregation type *j* and the true parental genotypes *m* and *n* is: 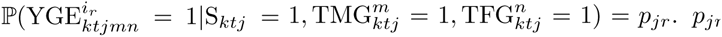 is equal to zero for homogametic individuals *r* = *hom* in any segregation type. *p_jr_* is also equal to zero for heterogametic individuals *r* = *het* in autosomal and X-hemizygous segregation types (*j* ≠ 2).

The probabilities of observing the parent and offspring genotypes can be defined when conditioned by all the hidden data of the model. The probability of observing 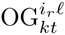, the genotype *ℓ* of offspring individual *i* of sex *r* at position *t* of contig *k*, given the segregation type *j*, the true parental genotypes *m* and *n*, the genotyping error *h* (either with an error *h* = *ε* or without error *h* = (1 – *ε*)) and the Y genotyping error *d* (either with an error *d* = *p_jr_* or without error *d* = (1 – *p_jr_*)) is 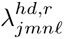. And similarly the probability of observing 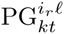, the genotype *ℓ* of parent individual *i* of sex *r* at position *t* of contig *k*, given all the hidden data is 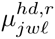, where *w* is the true genotype of the studied individual (either *m* or *n*). For instance, in the case of an autosomal segregation type (*j* = 1), if the heterogametic parent true genotype *n* is `AC` and the homogametic parent true genotype *m* is `AA` (as shown in Figure 1.a) and if no genotyping error has occurred (*h* = (1 – *ε*) and *d* = (1 – *p_jr_*)), then the probability 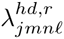 of observing genotype *ℓ* = `AA` in the offspring is 1/2 for both males and females. But in the case of a X/Y segregation type, as shown in Figure 1.b, then genotype `AC` is observed with probability 1 in males and genotype `AA` with probability 1 in females. Similarly, if the true homogametic parent genotype *m* is `AA` and no genotyping error has occurred, then the probability 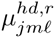 of observing genotype *ℓ* = `AA` in the homogametic parent is 1, and if a genotyping error occurred for this individual then all other genotypes *ℓ* = `AA` can be observed with probability 1/9 (as there are nine genotypes other than AA). In the case of an X/Y segregation type with the true heterogametic parent genotype *n* being `X`^A^`Y`^C^, the probability of observing genotype `AA` for this individual is 1 if there has been a Y genotyping error. All values for *λ* and *μ* can be found in Supplementary Table S1.

#### Inferences

An Expectation Maximisation (EM) algorithm is used to estimate the parameter values of the model *θ* = (*π, α, β, ε, p*). Detailed equations of the EM algorithm can be found in Supplementary text S1. Once parameters have been estimated, the *posterior* probabilities of the hidden data are computed given the observed data: Ŝ_*ktj*_ = ℙ(S_*ktj*_ |*G*) is the *posterior* probability of segregation type *j* at position *t* in contig *k*, given the observed data *G*. Using Bayes’ rule we get:

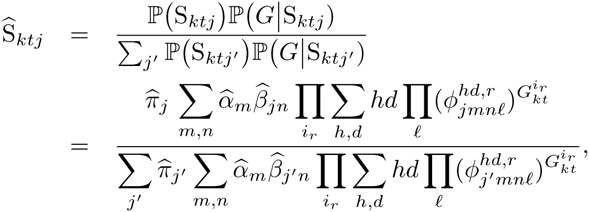

with *h* ∈ (*ε*̂, 1 – *ε*̂), *d* ∈ (*p*̂_*jr*_, 1 – *p*̂_*jr*_) and,

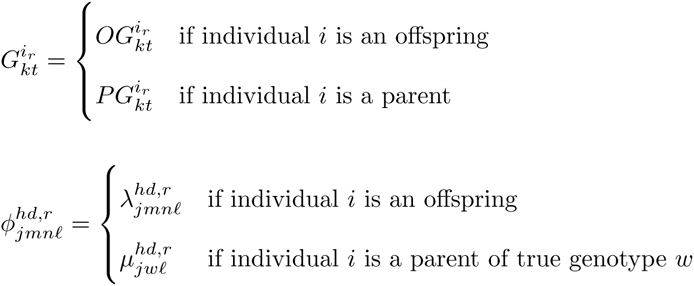

Similar derivations are done for the other posterior probabilities of hidden variables 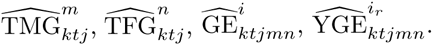.

Then, the segregation type of each position in the dataset can be inferred using the posterior segregation type Ŝ_*ktj*_. Contigs are attributed to a segregation category using positions that are polymorphic and informative. A position that was inferred as X or Z-hemizygous and that is polymorphic is always informative. A position that was inferred as autosomal or X/Y is considered informative only if the heterogametic parent is heterozygous and has a genotype that is different from the homogametic parent (otherwise it is not possible to differentiate between X/Y and autosomal segregation). The posterior segregation type of the contig is the average of the informative positions in the contig (assumed independent), with the positions being attributed a weight according to the posterior probability of genotyping errors (if a position has high genotyping error *posterior* probabilities it will be given less weight in the final decision for the contig segregation type),

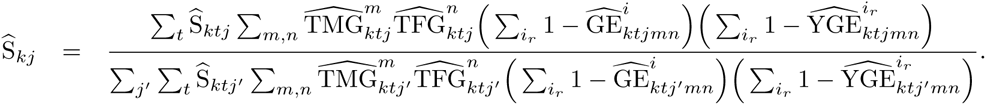

A contig is attributed to a sex-linked (X/Y or X-hemizygous) segregation type if its *posterior* probability of being X/Y plus X-hemizygous is higher than a tunable threshold (0.8 by default) and if the contig has at least one X/Y or X-hemizygous SNP without error (a genotyping error is inferred when its *posterior* probability is higher than 0.5). Similarly, a contig is attributed to an autosomal segregation if its *posterior* probability of being autosomal is higher than the chosen threshold and if the contig has at least one autosomal SNP without error. For each SNP the true parental genotypes are inferred as the ones that have the highest 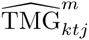 and 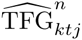 probabilities. Expression levels are retrieved using the X and Y (or Z and W) alleles predicted by this method and written in outputs. The model described above is adapted to X/Y and Z/W systems. Another version of the SEX-DETector model was written for U/V systems and can be found in Supplementary Text S2.

#### Bayesian Information Criterion (BIC) test for the presence of sex chromosomes

The Maximum Likelihood framework of the method allows the use of a model selection strategy to assess the presence of sex-linked genes in the dataset. A model *M* with the three possible segregation types can be compared to a model with only autosomal segregation type using the BIC, defined such that

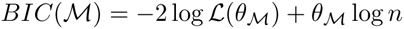

Where *BIC*(*M*) is the BIC value of model *M, L*(*θ_M_*) is the likelihood of the model, *θ_M_* is the number of free parameters of the model and n is the sample size. The model with the lower BIC value is chosen. It is also possible to test for a X/Y versus a Z/W system by comparing both BIC values. In case a model with sex chromosomes fits best the data but no sex-linked genes are inferred, then it means there are no sex chromosomes in the dataset. This could happen because of the extra Y genotyping error parameter *p*_2*het*_ that is specific to the model with sex chromosomes, which could account for mapping and genotyping errors in the data better than the genotyping error parameter *ε* alone.

### Data analysis

#### Plant material and sequencing

RNA-seq data were generated from a cross in *S. latifolia*, a dioecious plant that has sex chromosomes, and from a cross in *S. vulgaris*, a gynodioecious plant that does not have sex chromosomes. We used the following RNAseq libraries that were used in previous studies: Leuk144-3_father, a male from a wild population; U10_37_mother, a female from a ten-generation inbred line (Muyle, Zemp, et al., 2012); and their progeny (C1_01_male, C1_3_male, C1_04_male, C1_05_male, C1_26_female, C1_27_female, C1_29_female, C1_34_female). For *S. vulgaris* the hermaphrodite father came from a wild population (Guarda_1), the female mother from another wild population (Seebach_2) and their hermaphrodite (V1_1, V1_2, V1_4) and female (V1_5, V1_8, V1_9) progeny. Individuals were grown in a temperature-controlled greenhouse. The QiagenRNeasy Mini Plant extraction kit was used to extract total RNA two times separately from four flower buds at developmental stages B1–B2 after removing the calyx. Samples were treated additionally with QiagenDNase. RNA quality was assessed with an Aligent Bioanalyzer (RIN.9 larger than 9) and quantity with an Invitrogen Qubit. An intron-spanning PCR product was checked on an agarose gel to exclude the possibility of genomic DNA contamination. Then, the two extractions of the same individual were pooled. Individuals were tagged and then pooled for sequencing. Samples were sequenced by GATC, Konstanz, Germany on an Illumina HiSeq2000 following an Illumina paired-end protocol (fragment lengths 150-250bp, 100 bp sequenced from each end). A normalized 454 library was generated for *S. latifolia* using bud extracts from 4 different developmental stages.

#### Assembly

Adaptors, low quality and identical reads were removed. The transcriptome was then assembled using trinity (Haas et al., 2013) on the combined 10 individuals described previously as well as the 6 individuals from (Muyle, Zemp, et al., 2012) and the normalized 454 sequencing that was transformed to illumina using 454-to-illumina-transformed-reads (since trinity cannot take 454 reads as input). Then, isoforms were collapsed using /trinity-plugins/rsem-1.2.0/rsem-prepare-reference. PolyA tails and ribosomal RNAs were removed using ribopicker. ORFs were predicted with trinity transcript_t_bes_scorin_ORFs.pl. In order to increase the probability of assembling X and Y sequences in the same contig, ORFs were further assembled using CAP3 (cap3 - p 70, Version Date: 10/15/07, Huang and Madan, 1999) inside of trinity components.

#### Mapping, genotyping and segregation inference

Illumina reads from the 10 individuals of the cross were mapped onto the assembly using BWA (version 0.6.2, bwa aln - n 5 and bwa sampe, Li and Durbin, 2009). The libraries were then merged using SAMTOOLS (Version 0.1.18, Li, Handsaker, et al., 2009). The obtained alignments were locally realigned using IndelRealigner (GATK, DePristo et al., 2011; McKenna et al., 2010) and were analysed using reads2snp (Version 3.0, - fis 0 - model M2 - outpu_genotype best - multLalleles acc - mimcoverage 3 - par false, Tsagkogeorga et al., 2012) in order to genotype individuals at each loci while allowing for biases in allele expression and not cleaning for paralogous SNPs as X/Y SNPs tend to be filtered out by paraclean (the program that removes paralogous positions, Gayral et al., 2013). SEX-DETector was then used to infer contig segregation types after estimation of parameters using an EM algorithm. Posterior segregation types probabilities were filtered to be higher than 0.8. All these steps are implemented in a Galaxy workflow (see pipeline in Figure 2).

**Figure 2.**
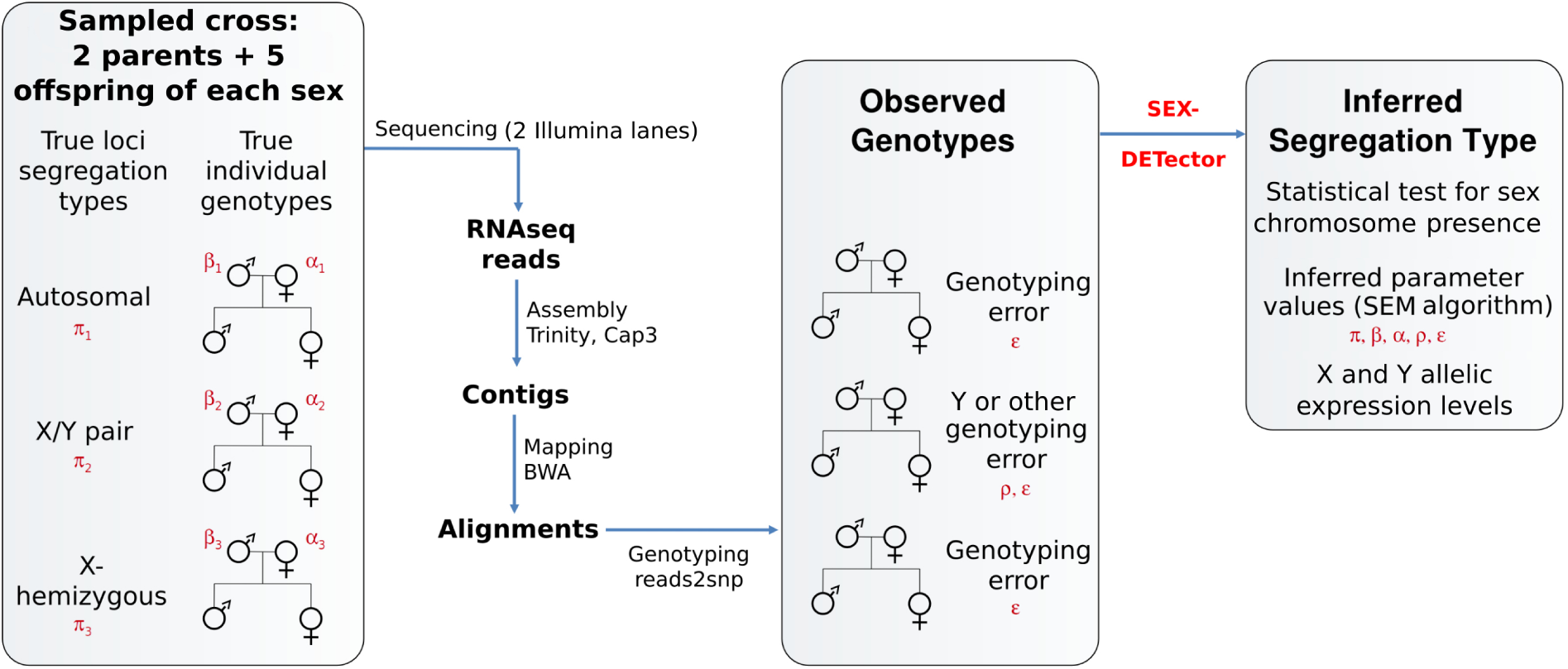
The SEX-DETector pipeline from required data to outputs. The advised number of offspring individual to sequence was determined with simulations. Parameters of the model are written in red: the segregation type *π*, the parents true genotype frequencies *α* and *β*, the genotyping error *ε* and the Y genotyping error *p*. An X/Y system is represented here but the pipeline is equivalent for a Z/W system. For a U/V system, only two individuals of each sex and one parent are advised and can be sequenced on a single Illumina Hi-seq 2000 lane. The pipeline was implemented in Galaxy.

#### The tester set, sensitivity and specificity

For various tests, we used 209 genes with previously known segregation type : 129 experimentally known autosomal genes, 31 experimentally known sex-linked genes (X/Y or X-hemizygous) and 49 X-linked CDS from BAC sequences (Supplementary Table S2). The sequences of these 209 genes were blasted (blast - e 1E-5, Altschul et al., 1990) onto the de novo assembly in order to find the corresponding ORF of each gene. Blasts were filtered for having a percentage of identity over 90% and an alignment length over 100bp and manually checked. Multiple RNA-seq contigs were accepted for a single gene if they matched different regions of the gene. If multiple contigs matched the same region of a gene, only the contig with the best identity percentage was kept. The gene was considered inferred as sex-linked if at least one of his matching contig was sex-linked. The inferred status of the genes by SEX-DETector was then used to compute specificity and sensitivity values. The same approach was used to compute sensitivity and specificity values for the three previous studies that inferred *S. latifolia* RNA-seq contig segregation patterns (Bergero and Charlesworth, 2011; Chibalina and Filatov, 2011; Muyle, Zemp, et al., 2012).

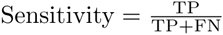

Sensitivity (or true positive rate) measures the capacity to detect true positives TP (genes that are sex-linked and inferred as such by the method). False negatives FN are sex-linked genes missed by the method.

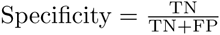

Specificity (or true negative rate) measures the capacity to avoid false positives FP (genes that are not sex-linked but inferred as such by the method). True negatives TN are non sex-linked genes detected as such by the method.

### Simulations

Simulated genotypes were used in order to test the effect of various parameters on the sensitivity and specificity of SEX-DETector. Sequences were first simulated for two parents (or a single parent in the case of a UV system) using the program ms to generate a coalescent tree (Hudson, 2002, see Supplementary Figure S1 and then the program seq-gen to generate sequences using the ms tree and molecular evolution parameters (version 1.3.2x, seq-gen - mHKY - l contig_length - f 0.26 0.21 0.23 0.3 - t 2 - s theta, Rambaut and Grassly, 1997). Different types of sequences were generated: either autosomal (ms 4 1 - T) or X/Y (ms 4 1 - T - I 2 3 1 - n 2 0.25 - n 1 0.75 - ej XY_divergenc_time 2 1 - eN XY_divergenc_time 1) or X-hemizygous (same parameters as X/Y but no Y sequence drawn) or U/V (ms 2 1 - T - I 2 1 1 - n 2 0.5 - n 1 0.5 - ej UV_divergence 2 1 - eN UV_divergence 1). Then, allele segregation was randomly carried on for a given number of progeny of each sex, using the segregation pattern determined when generating sequences with ms and seq-gen (see Supplementary Table S1 for segregation tables). *θ* = 4*N_e_μ* was set to 0.0275 as estimated in *S. latifolia* by (Qiu et al., 2010). *μ* was set to 10^−7^, which implies that 4*N_e_* was equal to ~70,000. Contig lengths were randomly assigned from the observed distribution of contig lengths of the *S. latifolia* assembly presented previously. Equilibrium frequencies used for seq-gen were retrieved from SEX-DETector inferences on the observed *S. latifolia* data. The transition to transversion ratio was set to 2 as inferred by PAML (Yang, 2007) on *S. latifolia* data (Kafer et al., 2013). The rate of genotyping error (*ε*) was set to 0.01 and the rate of Y genotyping error (*p*_2*het*_) was set to 0.13 as inferred by SEX-DETector on the observed *S. latifolia* data. Five types of datasets were simulated, with ten repetitions for each set of parameters and 10,000 contigs simulated for each dataset:

- Effect of X-Y divergence: Five different X-Y divergence times in units of 4*N_e_* generations were tested, either *S. latifolia* X-Y divergence time (4.5My) or 10 times or 100 times older or younger. The proportion of X-hemizygous contigs among sex-linked contigs was set accordingly to X-Y divergence time: 0.002, 0.02, 0.2, 0.6 and 1 for respectively 45,000 years, 450,000 years, 4.5 My, 45 My and 450 My divergence time. As well as the proportion of Y genotyping error (since Y expression is known to decrease with X-Y divergence): 0, 0.01, 0.13, 0.2 and 1 respectively. Four offspring of each sex were simulated. The proportion of sex-linked contigs was set to 10%.
- Effect of the number of sex-linked contigs: Five different proportions of sex-linked contigs (X/Y pairs or X hemizygous) were tested: 30% (3000 sex-linked contigs out of 10,000), 5%, 1%, 0,1% and 0,01%. Four offspring of each sex were simulated and X-Y divergence was set to 4.5 My.
- Effect of theta: Three different *θ* = 4*N_e_μ* (polymorphism) were tested: 0.000275, 0.00275 and 0.0275. Five offspring of each sex were simulated and X-Y divergence was set to 4.5 My, the X-Y divergence time in unit of 4*N_e_* generations varied accordingly to the value of theta. The proportion of sex-linked contigs was set to 10%.
- Effect of the number of individuals in Z/W and X/Y systems: Nine different numbers of offspring individuals of each sex were tested for the X/Y system: 2, 3, 4, 5, 6, 7, 8, 12 or 16 individuals of each sex. Sex chromosome size was set to 10% and X-Y/Z-W divergence to 4.5 My.
- Effect of the number of individuals in U/V systems: Eight different numbers of offspring individuals of each sex were tested for the U/V system: 1, 2, 3, 4, 5, 6, 7 or 8 individuals of each sex. Sex chromosome size was set to 10% and U-V divergence to 4.5 My.

For each simulated dataset, segregation types were inferred using SEX-DETector and were compared to the true segregation types in order to compute sensitivity and specificity values.

### Implementation and availability

The SEX-DETector code was written in perl and a Galaxy workflow was also developed (see user guide and source codes at http://lbbe.univ-lyon1.fr/-SEX-DETector-.html).

## Results

### The SEX-DETector pipeline

SEX-DETector takes as input file the genotypes of a cross (parents and progeny of each sex) for different contigs of an assembly. This data can typically be obtained from RNA-seq. The output is the inferred segregation type for every SNP and contig of the data (autosomal, X/Y or X-hemizygous, see Figure 1 for an example) along with allelic X and Y (or Z and W or U and V) expression levels. The SEX-DETector pipeline is pictured in Figure 2 and has been implemented as a Galaxy workflow. Simulations showed that the sequencing of two parents plus five progeny of each sex (12 individuals in total) is sufficient to obtain good results in an X/Y or Z/W system (see below). The use of RNA-seq lowers the cost, especially for species with large genomes. The pipeline can easily be modified to handle DNA-seq data. In order to obtain sufficient coverage, sequencing 20 to 25 million reads per individual is recommended (i.e. two Illumina lanes for twelve individuals on a Hiseq 2000). It is also recommended to use RNA extracted from a complex tissue where many genes would be expressed, especially sex-determining genes (e.g. flower buds in plants). The parents should be sampled from two different populations in order to increase the number of SNPs and therefore the power of the method. RNA-seq reads can be assembled into transcripts using Trinity (Haas et al., 2013). An important step after that is to further assemble transcripts, for example with Cap3 (Huang and Madan, 1999), in order to coassemble X and Y alleles in a single X/Y contig and avoid X/Y contigs to be misinferred as X-hemizygous. After mapping the reads onto the assembly, all individuals can be genotyped. The use of reads2snp (Tsagkogeorga et al., 2012) is highly recommended as it was designed for RNA-seq data in non-model organisms and allows for allelic expression biases, a key parameter when dealing with sex chromosomes and the poorly expressed Y alleles (Bachtrog, 2013). SEX-DETector takes reads2snp output as input.

SEX-DETector uses a probabilistic model to cluster contigs into segregation types. The parameters of the SEX-DETector model are estimated from the data using an EM algorithm (see Materials and Methods for details). Parameter *π*, for segregation type frequencies, allows to adapt to different species with different sex chromosome sizes. Parameters *α* and *β*, the parental genotype frequencies, accommodate for the heterozygosity level of the parents as well as the base composition of the species. The probability for a genotyping error to occur ε accounts for possible differences between observed and true genotypes (due to sequencing, mapping or genotyping errors). A specific Y genotyping error parameter *p* fits the high genotyping error rate for Y alleles due to degeneration and lower expression.

### Testing our pipeline’s performance using a *Silene latifolia* dataset

The SEX-DETector pipeline (Figure 2) was run on a cross dataset sequenced by RNA-seq in the plant Silene latifolia. This dioecious species was interesting for benchmarking our method/pipeline as its sex chromosomes are well known: they are relatively recent (~5 MY old, Rautenberg et al., 2010) but old enought to be clearly heteromorphic (the X is 400 Mb and the Y is 550 Mb, Matsunaga et al., 1994). X-Y synonymous divergence ranges from 5 to 25% (Bergero, Forrest, et al., 2007), *S. latifolia* thus represents a system of intermediate age. Also, a tester set of 209 genes for which segregation type has been established is available in this species (Supplementary Table S2). The dataset that was used here consists of a cross (two parents and four offspring of each sex). RNA-seq data were obtained for each of these individuals tagged separately and the reads were assembled using Trinity and then Cap3, the final assembly included 46,178 ORFs (Table 1). RNA-seq reads were mapped onto this assembly (see Supplementary Table S3 for library sizes and mapping statistics) and genotyping was done for each individual using reads2snp. SEX-DETector was run on the genotyping data to infer autosomal and sex-linked genes (Table 1). Figure 3 shows examples from the tester set. For some genes, all SNPs show clearly the same correct segregation type (Figure 3.A–C), whereas in some genes mixed segregation patterns were inferred, which we attribute to co-assembly of recent paralogs or other assembly/mapping problems (see Figure 3.D). These mixed cases can be filtered by the user, although they can happen in true sex-linked genes as is the case in Figure 3.D.

**Table 1:** Results of the SEX-DETector pipeline on the *S. latifolia* dataset.

| **ORF Types** | **Numbers** |
| --- | --- |
| ORFs in final assembly | 46178 |
| ORFs with enough coverage to be studied | 43901 |
| ORFs with enough informative SNPs to compute a segregation probability | 17189 |
| ORFs with posterior segregation probability over 0.8 | 15164 |
| ORFs assigned to an autosomal segregation type | 13807 (91%) |
| ORFs assigned to a X/Y segregation type | 1025 (7%) |
| ORFs assigned to a X hemizygous segregation type | 332 (2%) |

**Figure 3.**
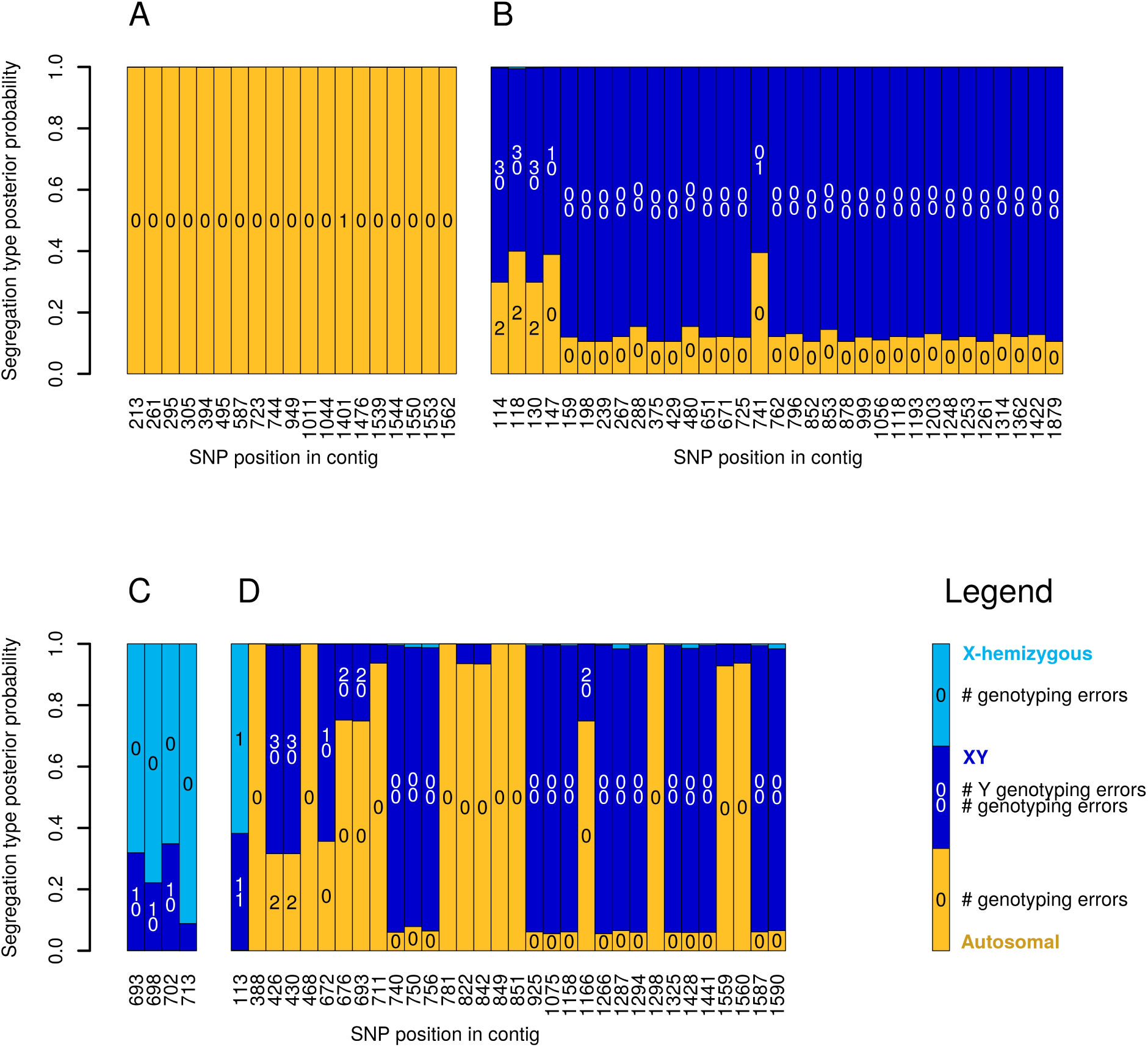
Results of the SEX-DETector pipeline for known S. *latifolia* genes. Segregation type posterior probabilities are shown for each informative SNP (see Materials and Methodss), see legend on figure for colour code, and inferred number of genotyping errors (see Materials and Methodss) are shown inside the bars. A) SlE72 is known to be autosomal, its weighted autosomal mean probability (see Materials and Methodss) is 0.99. B) SlCypX is known to be X/Y, its weighted sex-linked mean probability is 0.96. C) WUS1 is known to be X-hemizygous, its weighted sex-linked mean probability is 0.99. D) BAC284N5-CDS1_SlX6a is known to be sex-linked, its weighted sex-linked mean probability is 0.82.

We used our tester set to measure the performance of our pipeline, i.e. estimate its sensitivity (the capacity to detect true sex-linked genes) and specificity (the capacity not to assign autosomal genes as sex-linked, see Materials and Methodss). 83% of the known sex-linked genes expressed in the RNA-seq data used here (i.e. flower bud) were detected, indicating a high sensitivity. We obtained a specificity of 99% for this dataset as one gene, OxRZn, was supposedly wrongly assigned as a sex-linked gene by SEX-DETector. However, this gene was earlier assessed as autosomal on the basis of the absence of male specific alleles (Marais et al., 2011) and SEX-DETector assigned it to a sex-linked category because of two clear X-hemizygous SNPs, without genotyping error. It is therefore likely that OxRZn is in fact a true positive and more research on that gene is required.

### Comparing our pipeline to others using a *S. latifolia* dataset

We compared the performance of our pipeline to those used in previous work on inferring sex-linkage with RNA-seq data in *S. latifolia* (Bergero and Charlesworth, 2011; Chibalina and Filatov, 2011; Muyle, Zemp, et al., 2012). Those pipelines differ in many ways, and the data themselves can be different. In previous work, offspring individuals of the same sex were sometimes pooled before sequencing (Bergero and Charlesworth, 2011; Chibalina and Filatov, 2011). We used again the tester set of 209 *S. latifolia* genes with known segregation types, which we blasted onto each dataset to find the corresponding contigs and their inferred segregation type (for details see Supplementary Table S2). Because the different pipelines require different types of data (pooled progeny versus individually-tagged offspring) and with different read coverages, we computed sensitivity on all known genes (expressed or not). Our pipeline outperformed other pipelines in terms of sensitivity, while specificity was comparable (Table 2), which indicates that SEX-DETector can uncover more sex-linked contigs, without increasing the rate of false positives.

**Table 2:** Comparison with other methods: sensitivity and specificity values (see Materials and Methodss) obtained with different methods using 209 known *S. latifolia* genes.

|  | **Sensitivity** | **Specificity** |
| --- | --- | --- |
| **SEX-DETECTOR** | 0.625 (0.509 – 0.731) | 0.99 (0.958 – 0.999) |
| **Muyle, Zemp, et al., 2012** | 0.275 (0.181 – 0.386) | 0.984 (0.945 – 0.998) |
| **Bergero and Charlesworth, 2011** | 0.25 (0.159 – 0.359) | 0.992 (0.958 – 0.999) |
| **Chibalina and Filatov, 2011** | 0.425 (0.315 – 0.541) | 1 (0.972 – 1) |

As further analysis showed, this was due to overly conservative filtering in previous work. To exclude false positives, genes with at least 5 sex-linked SNPs were retained in previous studies. More filtering was done by excluding contigs with autosomal SNPs (Bergero and Charlesworth, 2011; Hough et al., 2014). As shown in Figure 4, keeping only contigs with at least 5 sex-linked SNPs removes nearly half of the contigs inferred as sex-linked by SEX-DETector, many of which have a high posterior probability. Excluding further those with autosomal SNPs (keeping those with sex-linked SNPs only) removes 74% of the contigs (Figre 4.B). Comparatively, SEX-DETector removes 12% of contigs when filtering for a posterior probability higher than 0.8 (Table 1), as most genes have a very high posterior segregation type probability which indicates a strong signal in the data and illustrates the benefits of using a model-based approach.

**Figure 4.**
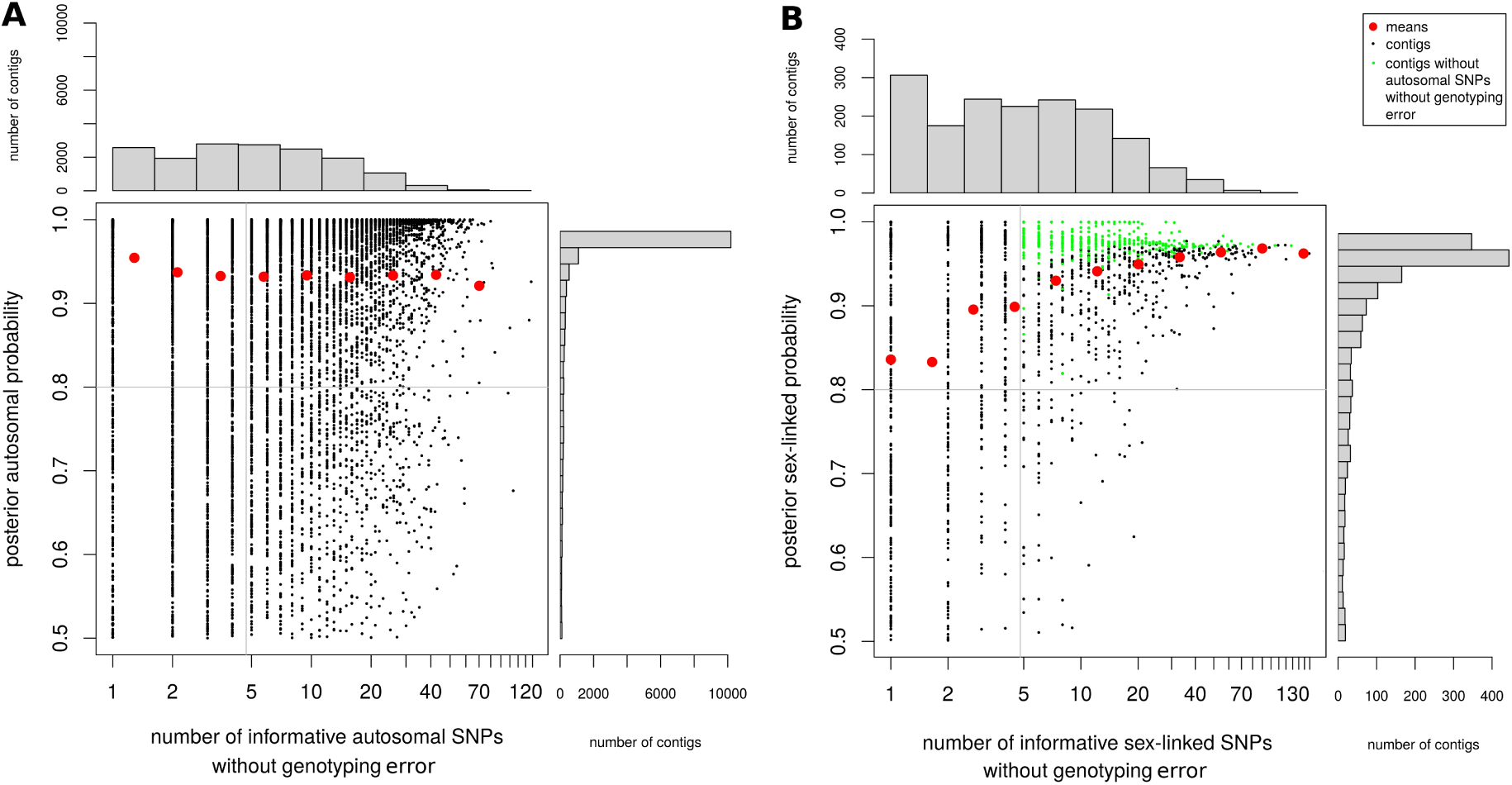
Performance of the method. The number of SNPs without genotyping error was plotted against the posterior segregation type probability for each autosomal (A) and sex-linked (B) contigs of the *S. latifolia* dataset. The distributions of both variables are shown, and means for each category on the histograms are indicated by red dots. Sex-linked genes that remain after filters that are commonly applied in empirical methods are shown in green (at least five sex-linked SNPs and no autosomal SNPs). SEX-DETector on the other hand filters for a posterior probability above 0.8 (horizontal line on graph) and at leat one sex-linked SNP, so that more contigs can inferred as sex-linked, without increasing the false positive rate compared to other empirical methods (Table 2).

### Simulations show that SEX-DETector requires a modest experimental effort and works on different sex chromosome systems

We simulated genotypes for a cross (parents and progeny) by generating coalescent trees with either autosomal or sex-linked history (Supplementary Figure S1) and generated the parental sequences using these trees and molecular evolution parameters. Progeny genotypes were obtained by random segregation of alleles from the parents and a genotyping error layer was added (see Materials and Methodss). 10,000 contigs were simulated for each dataset.

In order to know how many offspring of each sex should be sequenced to achieve the best sensitivity and specificity trade-off using SEX-DETector, we varied the number of progeny individuals in the simulations. For an X/Y or Z/W system, optimal results were obtained when sequencing five progeny individuals of each sex (Figure 5.A); sequencing more progeny individuals did not improve the results further. This suggests that sequencing 12 individuals (two parents and five progeny individuals of each sex) may be sufficient to achieve optimal performances with SEX-DETector on an X/Y or Z/W system. For a U/V system, two progeny individuals of each sex seems sufficient to obtain optimal SEX-DETector performance (Figure 5.B), which suggests that sequencing five individuals (the sporophyte parent and two progeny of each sex) may be enough in the case of a U/V system. Our simulations thus suggest that SEX-DETector requires a modest experimental effort to reliably identify expressed sex-linked genes.

**Figure 5.**
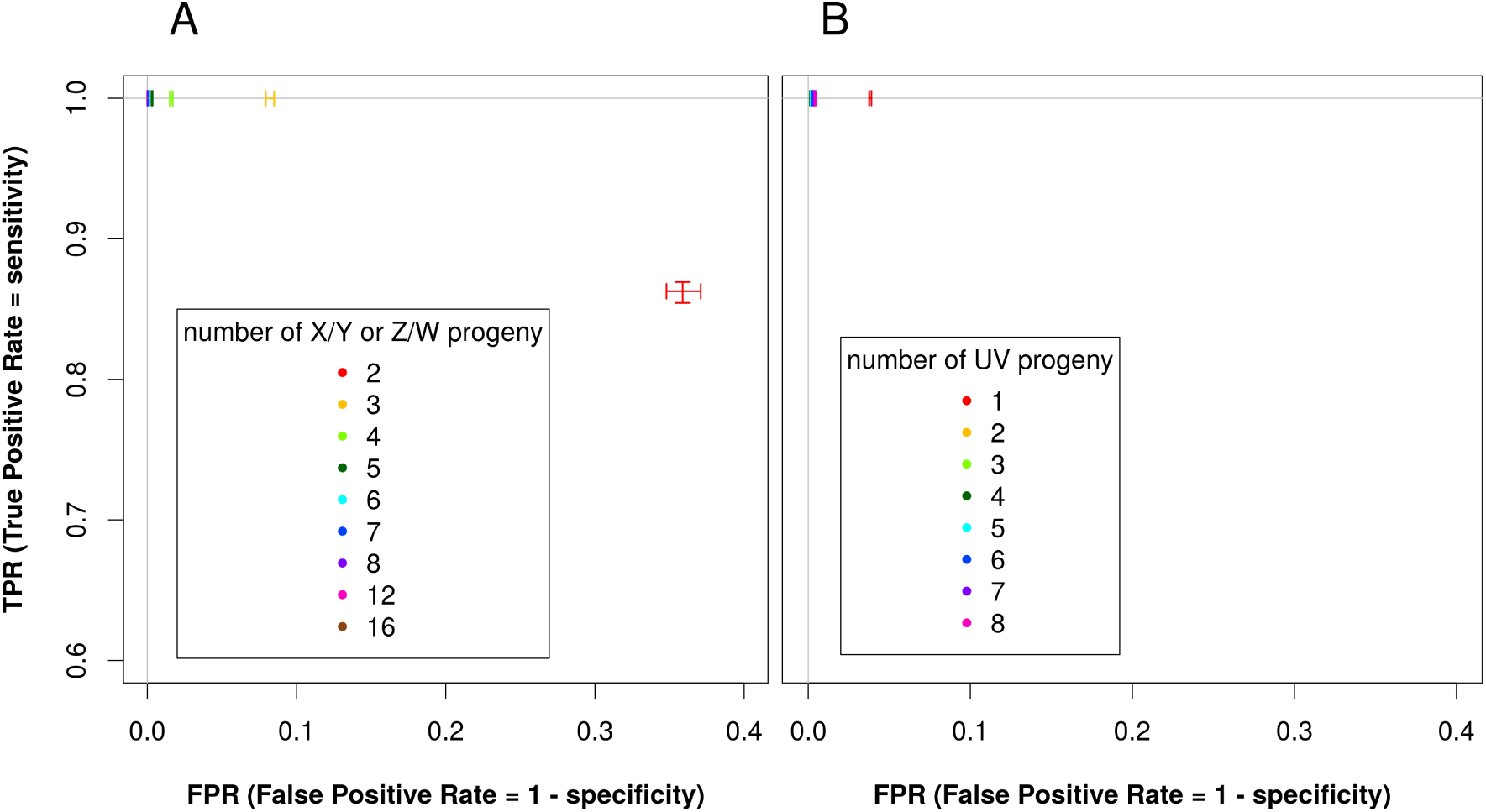
Measure of the effect of the number of sequenced offspring individuals using simulations. ROC curve (true positive rate represented as a function of false positive rate), showing the effect of the number of progeny sequenced on sensitivity (TPR, true positive rate) and specificity (1-FPR, false positive rate) in simulated data. A perfect classification of contigs would lead to a point having TPR equal to one and FPR equal to zero (top left corner of the graph). **A**) X/Y or Z/W sex determination system (all points overlap in the top left corner when over five progeny of each sex are used). **B**) U/V system (all points overlap in the top left corner when over two progeny of each sex are used).

In order to assess the applicability of SEX-DETector to different types of sex chromosomes (old versus young, homomorphic versus heteromorphic) and species (highly versus weakly polymorphic), we used the same simulation procedure and tested the effect of one parameter at a time on SEX-DETector sensitivity and specificity. In our simulations, the degree of polymorphism within species had no influence on the performance of our method (Supplementary Figure S2.A). As for the influence of the size of the nonrecombining region (homomorphic or heteromorphic sex chromosomes), it was tested using different % of sex-linked genes in a genome with no effect on the performance of SEX-DETector (Supplementary Figure S2.B). The limit of detection of a sex-linked contig was reached only when 1 sex-linked contig out of 10,000 contigs was present. Finally, the simulations indicated that our method is robust to X-Y divergence, as young and old sex chromosomes were evenly detected (Supplementary Figure S2.C).

### SEX-DETector identifies unknown sex chromosomes using model selection

It is common that the sex determination system is unknown in species with separated sexes, i.e. it is unknown whether they have sex chromosomes and if they do, what the system is (Z/W or X/Y). The likelihood-based framework of SEX-DETector allows us to test for these assumptions by comparing the model fit to the data using the Bayesian Information Criterion (BIC, see Materials and Methodss). In species for which sex determination is unknown, it is possible to compare models with and without sex chromosomes, and, if sex chromosomes are detected, it is possible to compare models with X/Y or Z/W system. This model selection strategy was tested on empirical and simulated data.

In the *S. latifolia* dataset, the best model inferred by SEX-DETector was a model with sex chromosomes as expected, with 1357 sex-linked contigs (which represents 9% of the contigs with a posterior probability higher than 0.8). In the *Silene vulgaris* dataset (a species without sex chromosomes), no sex-linked contigs were inferred, the best model fit to the data was thus a model without sex chromosomes as expected (see Materials and Methodss).

In order to know from which proportion of sex-linked genes sex chromosomes can be detected, we compared models on simulated data with varying numbers of sex-linked contigs out of 10,000 simulated contigs (Table 3 and Supplementary Table S4). When no sex-linked contigs were simulated, as expected the best model was the one without sex chromosomes. This was also the case when a single sex-linked contig was simulated. In this case, SEX-DETector could not detect it due to lack of information in the dataset. When ten or more sex-linked contigs were simulated, the best model was the one with sex chromosomes as expected. Thus, ten sex-linked contigs out of 10,000 provide sufficient information for SEX-DETector (i.e. 1 sex-linked gene out of 1000 genes can be detected).

**Table 3:** Model comparison using SEX-DETector on empirical datasets (*S. latifolia* (with sex chromosomes) and *S. vulgaris* (without sex chromosomes) and simulated X/Y datasets with varying number of sex-linked contigs out of 10,000 simulated contigs. The best model is chosen as the one having the lowest BIC value (see Materials and Methodss and Supplementary Table S4 for details).

|  |  | Best model | number of sex-linked genes in the best model |
| --- | --- | --- | --- |
| Empirical datasets | *Silene latifolia* (X/Y system) | X/Y | 1357 |
|  | *Silene vulgaris* (no sex chromosomes) | Z/W | 0 |
| Simulated datasets of 10,000 genes with different numbers of sex-linked genes (XY system) | 0 sex-linked genes | no sex chromosomes | 0 |
|  | 1 sex-linked genes | no sex chromosomes | 0 |
|  | 10 sex-linked genes | X/Y | 16–57 |
|  | 100 sex-linked genes | X/Y | 156–181 |
|  | 500 sex-linked genes | X/Y | 592–624 |
|  | 3000 sex-linked genes | X/Y | 3159–3200 |

Once the presence of sex chromosomes has been inferred, it can be tested whether the system is X/Y or Z/W. The model comparison between X/Y and Z/W systems worked on both empirical and simulated data: the best model for *S. latifolia* was, as expected, the X/Y system (Table 3 and Supplementary Table S4).

## Discussion

To summarise, SEX-DETector implements a probabilistic model that is used to compute the posterior probabilities of being autosomal, X/Y and X-hemizygous (X-linked copy only) for each RNA-seq contig in data from a full-sib family. The method is suitable for any sex chromosome type (XY, ZW, UV). SEX-DETector uses genotypes obtained from a genotyper specifically designed for RNA-seq data (Gayral et al., 2013; Tsagkogeorga et al., 2012). This genotyper takes unequal allelic expression into account, which is particularly important as Y (or W) copies tend to be less expressed than their X (or Z) counterpart (reviewed in Bachtrog, 2013). The SEX-DETector model also accounts for genotyping errors. The pipeline that includes steps from assembly to sex-linkage inference (Figure 2) was implemented in Galaxy for easier use. The pipeline was successfully tested on RNA-seq data of a family from *Silene latifolia*, a dioecious plant with relatively recent but heteromorphic sex chromosomes. Genes have previously been experimentally characterised as autosomal or sex-linked in this species, which made it possible to assess the performances of the method. 83% of known sex-linked genes that were expressed in the sampled tissue could be identified using the SEX-DETector pipeline. Sensitivity and specificity values were used to compare SEX-DETector to other RNA-seq based approaches that used *S. latifolia* datasets (Bergero and Charlesworth, 2011; Chibalina and Filatov, 2011; Muyle, Zemp, et al., 2012). SEX-DETector showed a much higher sensitivity (0.63 compared to 0.25-0.43) while specificity remained close to 1. Thanks to a statistically grounded method, the SEX-DETector pipeline can detect many more genes than previous approaches, while keeping the inferences extremely reliable. The SEX-DETector pipeline was also run on a comparable RNA-seq data from Silene vulgaris (a plant without sex chromosomes), and yielded no sex-linked genes, as expected. We further tested the SEX-DETector method using simulations, which indicated good performance on different sex chromosome systems (old or young and homomorphic or heteromorphic). Simulations also showed that few individuals need to be sequenced for optimal results (under 12 individuals for a ZW or XY system and five individuals for a UV system). This makes the strategy very accessible given the cost of RNA-seq, in particular in specie with large genomes. The likelihood framework of SEX-DETector makes it possible to assess the presence and type of sex chromosomes in the data using a model comparison strategy. This procedure proved efficient on empirical and simulated data, as long as more than 1 gene in 10,000 was sex-linked in the data.

The downside of using RNA-seq data is, of course, that only expressed genes can be identified by the SEX-DETector pipeline. This can be overcome by the use of DNA-seq data, or the combination of multiple tissues for RNA-seq data. Moreover, because Y-specific genes cannot be differentiated from autosomal male-specific genes in RNA-seq data, Y genes are not inferred by SEX-DETector unless they coassemble with an X counterpart. This requirement makes the method less adapted to old sex chromosome systems where X and Y copies of a given gene could be too diverged to coassemble. However, X copies can still be identified on their own if the Y copy is absent or did not coassemble with the X (Figure 1.c). To try and identify missed Y contigs, X-hemizygous genes can be blasted onto male-specific contigs, which may represent the diverged Y copy. This was done for the 332 inferred X-hemizygous genes in the *S. latifolia* dataset, and only 5 of them had a significant match with a male-specific contig. This suggests that very few true X/Y gene pairs were wrongly inferred as X-hemizygous due to a too divergent Y. In *S. latifolia*, X-Y synonymous divergence ranges from 5% to 25% (Bergero, Forrest, et al., 2007). This is comparable to regions in human sex chromosomes that stopped recombining last: the mean X-Y synonymous divergence for strata 3, 4 and 5 in humans is respectively 30%, 10% and 5% (Skaletsky et al., 2003). SEX-DETector will therefore perform best in species with sex chromosomes of young or intermediate age, but will also work on recent strata of old systems. A complete list of cases where inferences could be difficult is provided in the online SEX-DETector user manual (p 3-4), along with cases where SEX-DETector could be applied to detect dominant loci associated with a phenotype, other than sex chromosomes.

Other approaches to identify sequences of sex chromosomes based on female and male DNA-seq data comparison can only detect regions where the X and the Y are divergent enough not to coassemble nor map onto one another (Akagi et al., 2014; Carvalho and Clark, 2013; Cortez et al., 2014; Vicoso and Bachtrog, 2011; Vicoso, Emerson, et al., 2013; Vicoso, Kaiser, et al., 2013). These approaches are therefore best suited to old sex chromosome systems. Other methods work on young systems but rely on genome sequencing (Al-Dous et al., 2011; Hou et al., 2015; Picq et al., 2014). Obtaining a reference genome can be difficult for non-model organisms, especially those with large genomes. In such cases, RNA-seq data is a lot cheaper. Therefore, SEX-DETector is a highly promising method for uncovering sex-chromosomes in non-model organisms, and especially those with young sex chromosomes. This type of sex chromosomes is expected in thousands of yet unstudied independent taxa across plants and animals (see Introduction and Bachtrog, Mank, et al., 2014; Ming et al., 2011; Renner, 2014), and probably many more in all eukaryotes.

## Acknowledgements

We thank Alex Widmer for access to the RNA-seq datasets and comments on the manuscript, Nicolas Galtier and Sylvain Glemin (ISEM-Montpellier) for useful discussions about reads2snp, Vincent Miele (LBBE) for SEX-DETector profiling and advice on code performance, Khalid Belkhir (ISEM-montpellier) for providing and adapting Galaxy wrappers for analyses used upstream of SEX-DETector (BWA, reads2snp) and Philippe Veber (LBBE) for help with Galaxy.

## Supplementary Material

**Supplementary Figure S1:**
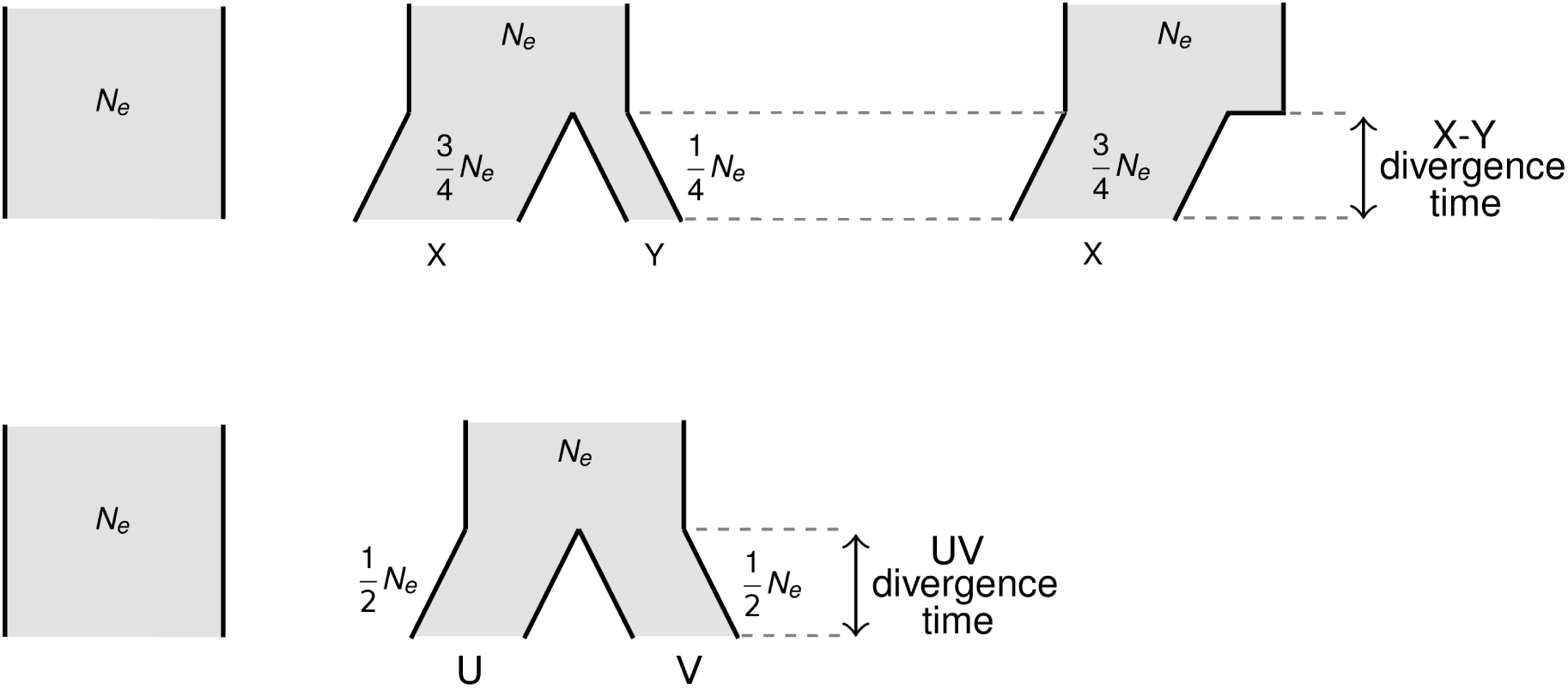
simulations design for the program ms (Hudson 2002), for X/Y system (upper part) and U/V system (lower part).

**Supplementary Figure S2:**
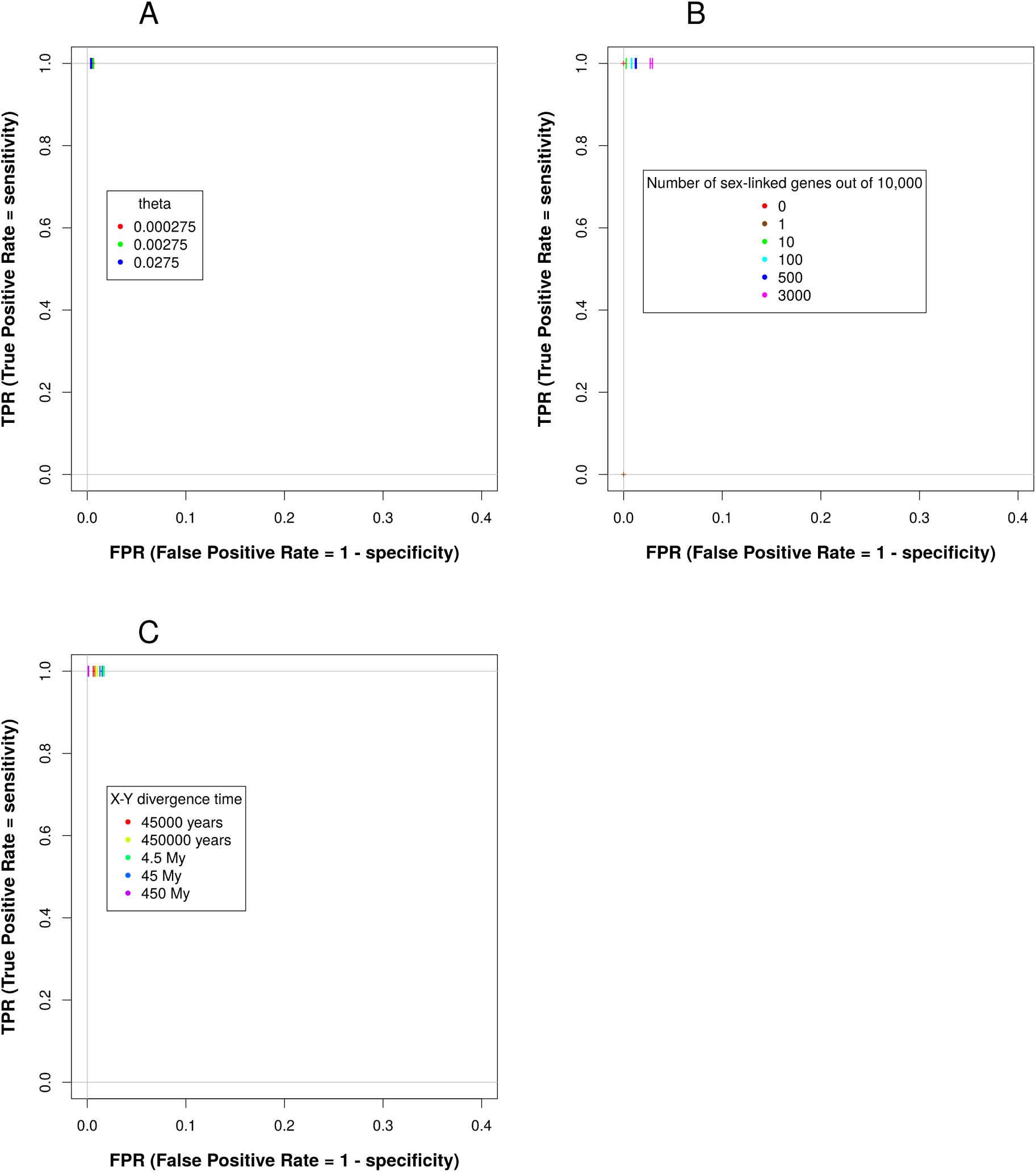
ROC curve showing the effect of different parameters on sensitivity (TPR, true positive rate) and specificity (1-FPR, false positive rate) in simulated data. A perfect classification of contigs would lead to a point having TPR equal to one and FPR equal to zero (top left corner of the graph). A) effect of X-Y divergence time. B) effect of the number of sex-linked contigs out of 10,000 simulated contigs. C) effect of theta (polymorphism).

**Supplementary Text S1:** Detailed explanation of the model for X/Y and Z/W systems.

**Supplementary Text S2:** Detailed explanation of the model for U/V systems.

**Supplementary Table S1:** Segregation tables, observed genotypes probabilities given true parent genotypes, segregation types and genotyping errors.

**Supplementary Table S2:** Known genes used to test SEX-DETector and compare it with other methods in *S. latifolia*.

**Supplementary Table S3:** Library sizes (in number of reads) and mapping statistics.

**Supplementary Table S4:** Details of model comparison using SEX-DETector on empirical datasets (Silene latifolia which has sex chromosomes and Silene vulgaris which does not have sex chromosomes) and simulated X/Y datasets with varying number of sex-linked contigs out of 10,000 simulated contigs. The best model is chosen as the one having the lowest BIC value (bold and stressed).

